# Monkey V1 epidural field potentials provide significant information about stimulus location, size, shape, and color

**DOI:** 10.1101/2020.12.29.424571

**Authors:** Benjamin Fischer, Detlef Wegener

## Abstract

Brain signal recordings with epidural microarrays constitute a low-invasive approach for recording distributed neuronal signals. Epidural field potentials (EFPs) may serve as a safe and highly beneficial signal source for a variety of research questions arising from both basic and applied neuroscience. A wider use of these signals, however, is constrained by a lack of data on their specific information content. Here, we make use of the high spatial resolution and the columnar organization of macaque primary visual cortex (V1) to investigate whether and to what extent EFP signals preserve information about various visual stimulus features. Two monkeys were presented with different feature combinations of location, size, shape, and color, yielding a total of 375 stimulus conditions. Visual features were chosen to access different spatial levels of functional organization. We found that, besides being highly specific for locational information, EFPs were significantly modulated by small differences in size, shape, and color, allowing for high stimulus classification rates even at the single-trial level. The results support the notion that EFPs constitute a low-invasive, highly beneficial signal source for longer-term recordings for medical and basic research by showing that they convey significant and reliable information about constituent features of activating stimuli.

## Introduction

Multielectrode arrays placed on top of the brain provide brain signals for studying dynamic neuronal interactions over large distances, within and across distinct brain areas^1,2,3^. They also offer new possibilities for future therapeutic purposes and brain-computer interfacing. Epidural multielectrode arrays, in particular, provide a low-invasive approach for recording brain activity with higher signal quality and less vulnerability to motor and ocular artifacts than electroencephalography^4,5^. On the other hand, epidural field potentials (EFPs) have presumably lower specificity than intracortically recorded signals, primarily due to the larger distance between electrodes and neurons and larger cortical spread of the signal^6^. Compared to signals from subdurally implanted microelectrode arrays, EFPs were found to have a higher noise level and to be more difficult to decode by some studies^7,8^. Other work, in contrast, reported consistent decoding performance between EFP and local field potential (LFP) and no detrimental effects of the dura on signal feature detection^9,10^. These partly contradicting results leave some uncertainty regarding EFP selectivity and presumably constrain the choice of epidural recordings over more invasive approaches.

The great potential of EFPs for clinical and BCI-usage was recently emphasized by the finding that spatial information of V1 EFPs can be decoded from very short single trial fragments with a detection rate of consistently more than 90%^11^. In that study, objects were shown at several close-by locations; they consisted, however, of differently colored letter stimuli, such that decoding performance also relied on differences in stimulus size, orientation statistics, and color. Currently available data on visually evoked EFPs do not allow to disentangle the contribution of different visual features. More detailed knowledge on this is mandatory for their use in brain-computer interfacing, and highly important for basic research and clinical settings. The present study, therefore, attempts to investigate the information content of epidural signals in more detail, by recording EFPs from macaque primary visual cortex (V1) in response to different, spatial and non-spatial visual features.

Because location, size, orientation, and color address different topographic maps, analysis of their representation allows insights into the specificity of EFPs with respect to cortical functional architecture at different scales. First, V1 is retinotopically organized, with average epidural receptive field (ERF) size diameters of about 2.5 to 4-degree^11^. This is a factor of approximately 1.5 to 2.5 compared to both the LFP and multi-unit activity mapped with the same technique^12^. We address the question whether and to what extent stimulus location and stimulus size is represented in EFPs if differences in location and size are kept within, or even below, the range of ERF diameters. Second, orientation- and color-selective V1 neurons are organized in cortical columns and blobs, respectively, which (together with ocular dominance columns) built a cortical hypercolumn^13,14,15^. Hypercolumns span about one to two degrees of the parafoveal visual field^13^, hence the area represented by an ERF is likely to cover, at least partly, several hypercolumns. While this suggests that the EFP at a given electrode is integrating over multiple sets of orientation columns and color blobs, parts of a stimulus may activate distinct neuronal subsets and thus may shape the integrated response in a specific manner. We address the question whether and to what extent EFPs are modulated by the orientation statistics of a visual object (that is, its shape), and its color. The results of the study show that, albeit the EFP is a mass signal that integrates over presumably tens of thousands of neurons, it preserves significant information not only about spatial but non-spatial stimulus features, which can be decoded with high classification rates on the single-trial level. This renders the EFP a highly informative yet minimally invasive brain signal for chronic, long-term applications in both clinical settings and basic research.

## Results

We implanted two monkeys with an epidural microelectrode array^16^ over the left V1 hemisphere. Data were obtained from a total of 176 and 137 electrodes in Monkey (M) 1 and 2, respectively. ERFs were mainly located in the lower visual quadrant and were mapped by an automated bar-mapping procedure based on reverse correlation and geometric means of differently oriented bar trajectories^11,12^. Mean ERF diameter (defined as closed areas of ≥ 1 *Z*-score activation and size of ≥ 1 square-degree) was 3.0 ± SD 0.46 visual degree in M1 and 2.8-degree ± 0.66 in M2, recalculated from actual ERF area by assuming circular shape. Visual stimuli were placed at five locations within the lower right quadrant of the visual field, at 3.5, 5.8, and 8.2-degree eccentricity (Fig. 1a). Stimulus locations were fixed throughout the experiment and were not adjusted to ERF coordinates, such that stimulus coverage of individual ERFs varied. For the example shown in Fig. 1a, the ERF had most overlap with stimulus location 4 and minor overlap with stimulus location 1, with the ERF center located between the two. At each location, stimuli were shown in three different sizes (1, 1.2, and 1.4-degree in diameter, exemplified from inner to outer stimulus locations in Fig. 1a). Stimuli consisted of two triangles, two squares, and a circle, with triangle and square variants differing only in global orientation (Fig. 1b). All objects were shown in five different (four chromatic and one achromatic) equiluminant colors (Fig. 1b). All objects were chosen to exactly fit into an orbit of the specified size diameter and, per size, were matched for their number of pixels. Per trial, three stimuli were presented during a passive fixation task (Fig. 1c) in pseudo-random order.

**Figure 1.**
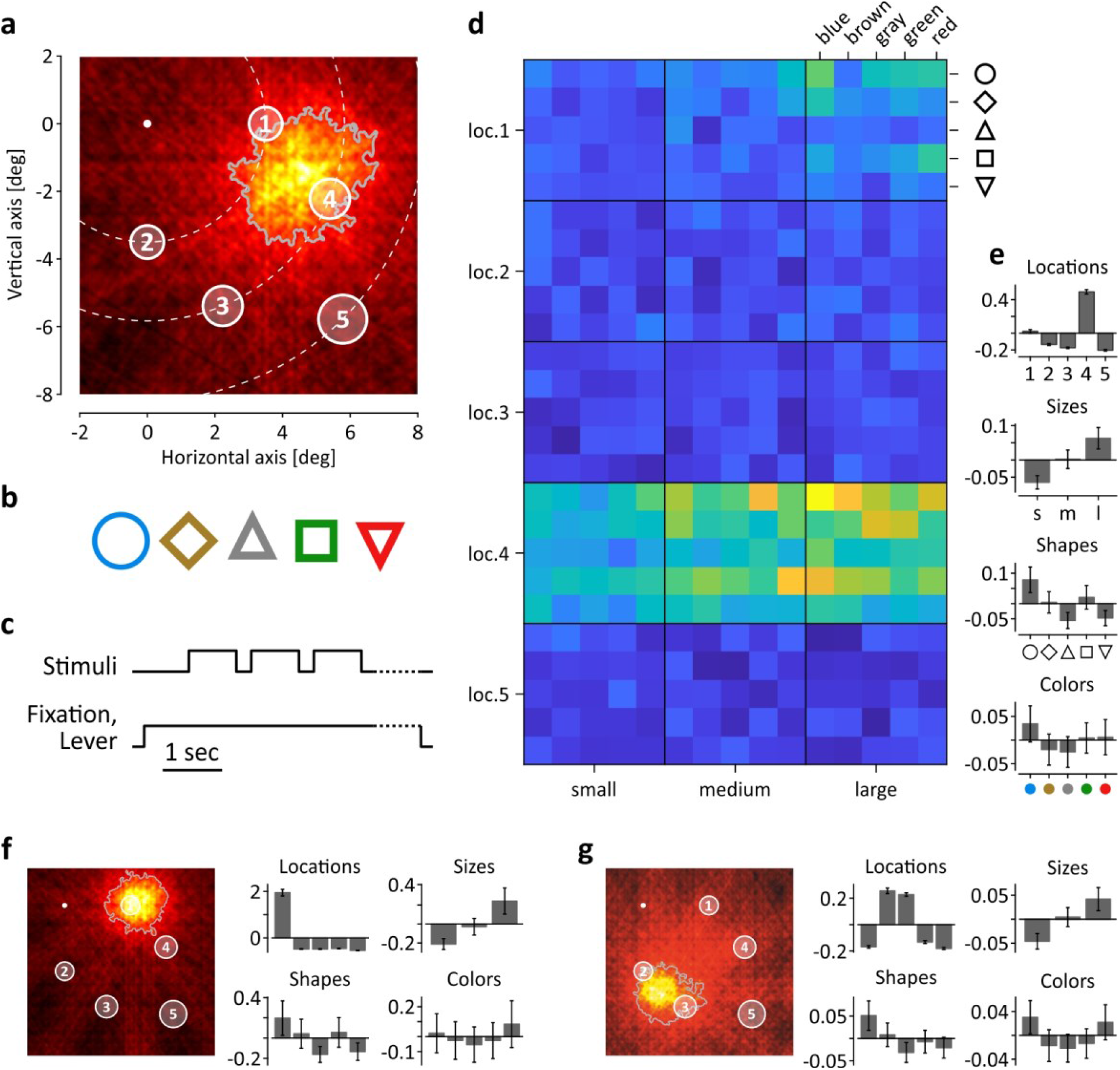
Visual stimulation, experimental paradigm, and broadband-gamma [25 – 160 Hz] responses at three example channels. (**a**) Response map illustrating ERF (area defined by gray line) and stimulus locations (open circles) in visual field coordinates. Increasing circle diameter from inner to outer locations represent the three different stimulus sizes. White dot indicates fixation point, dashed lines indicate iso-eccentricity at 3.5, 5.8, and 8.2 visual degree (deg). (**b**) Stimulus shapes and colors. Shape outlines were matched in thickness to provide identical number of pixels across shapes. Color isoluminance was matched by photometry. Each shape was shown at each location, in each color and size, providing a total of 375 stimulus conditions. (**c**) Time course of visual stimulation and behavioral paradigm. Dashed segment towards end of trial indicates randomized period of up to 1.1 sec before the to-be-detected event (dimming of fixation point). (**d**) Response matrix indicating relative mean broadband-gamma power for each of the 375 stimulus conditions. Warmer colors indicate higher power. (**e**) Barplots summarizing gamma-band responses (y-axis) per category and condition. For better visualization, mean power over all conditions was subtracted from each individual condition. Labelling of x-axis for sizes corresponds to small (s), medium (m), and large (l). (**f** - **g**) ERF maps and barplots for condition-specific differences in mean gamma power per category of two more example channels (f, M2; g, M1). Scaling of ERF maps corresponds to (a), x-axis of histograms corresponds to (d). Error lines in barplots indicate SEM throughout.

Figure 1d shows a response matrix of the results obtained from the channel shown in Fig. 1a for the onset (26 – 175 ms) broadband power (25 – 160 Hz) of the wavelet-transformed EFP, averaged over all trials (range over both monkeys: 21 − 39) of each of the 375 (5 locations * 3 sizes * 5 shapes * 5 colors) stimulus conditions. Neuronal gamma-band responses at this electrode were mainly elicited by stimulation at location 4, and to some extent at location 1, as to be expected from the ERF coordinates. Regarding stimulus size, responses were strongest for large stimuli and weakest for small ones, likely reflecting better overlap with the ERF and closer proximity to the ERF center with increasing stimulus size. Furthermore, activation at the electrode seemed to co-vary with object geometry but not with global orientation, and was stronger for blue stimuli than for other colors. For better illustration, Fig. 1e shows the normalized mean gamma power per feature category and condition. Results for e.g. the category size are shown as the difference between the mean over the trials available for small, medium, and large stimuli (irrespective of their location, shape, and color) and the overall mean. If not mentioned otherwise, all analyses that follow are based on this way of assignment and include the entire number of trials sorted by the conditions (e.g. blue, red) of the category (e.g. color) to be analyzed. Two more examples show a channel with ERF center coordinates closely matching stimulus location 1 (Fig. 1f) and another one having an ERF in-between two stimuli, with only moderate overlap to each of them (Fig. 1g). Notably, despite these differences in stimulus-ERF overlap, both channels, albeit taken from different animals, show gamma responses closely resembling the pattern of the former example regarding stimulus size and shape, and a slight bias towards red and blue stimulus color.

For a comprehensive analysis of feature-specific EFP modulations, we first determined the most-informative time-frequency range for each of the feature categories, as different stimulus features may activate distinct time-frequency ranges in V1^17,18,19^. Most-informative time-frequency ranges were estimated by an approach based on receiver-operating characteristics (ROC), as recently introduced^11^. Per monkey and category, the procedure delivers a time-frequency map indicating the range with best stimulus separability over all electrodes. During the initial stimulus onset response, location-specific modulation was most consistent in the high-gamma frequency range, while size- and shape-specific modulation was found in the low-to-mid-gamma range. Color-specific modulation was absent in the early onset response but was obvious during later response periods. Specific time-frequency ranges per monkey are indicated in Fig. 2. For each monkey, electrode, and feature category, analysis was based on the baseline-corrected wavelet power *WP* in the respective ROC-selected time-frequency range, averaged over all trials of a given stimulus condition.

**Figure 2.**
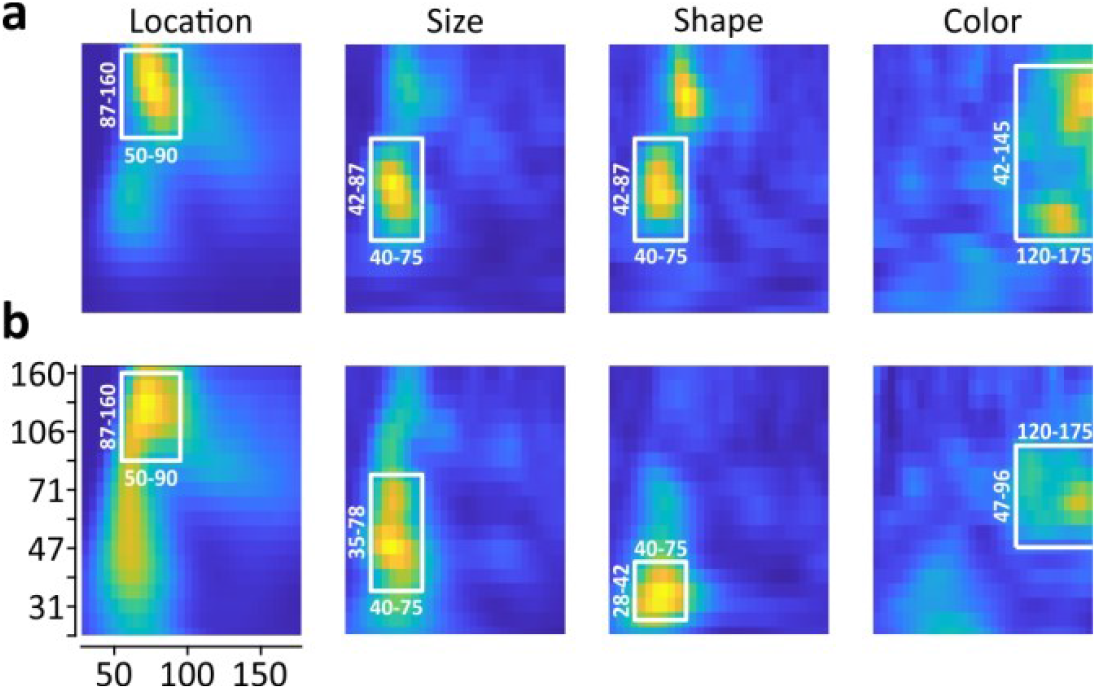
Time-frequency ranges used for EFP analysis. (**a** - **b**) Color maps show variance in area-under-the ROC curve over all electrodes, for stimuli sorted according to location, size, shape, and color, separately for M1 (a) and M2 (b). Y-axis indicates wavelet frequency ranges [Hz], x-axis indicates time [ms], aligned to stimulus onset. Whites boxes delineate the selected time-frequency range per feature and monkey. Color scaling is identical for all panels.

Results for the two monkeys were found to be very similar and provided matching statistical conclusions. For better overview, figures in the following sub-chapters summarize the results of both monkeys.

### Spatial sensitivity

Stimulus locations were chosen to cover different eccentricities and to differently overlap with ERFs to allow for analysis of the spatial spread of the signal and of feature selectivity as function of ERF-stimulus distance. Electrodes were assigned to the stimulus location with smallest Euclidian distance to their ERF center. Responses at 311 of 313 electrodes were found to be significantly dependent on stimulus location (Kruskal-Wallis test, all *P* < 0.028). 58 and 75% of electrodes in M1 and M2, respectively, showed a response to stimuli at their assigned location that was significantly different from the response to stimuli at any other location (pairwise tests with Tukey-Kramer correction for multiple comparison, all *P* < 0.05). Spatial sensitivity was, however, highly variable in magnitude (Fig. 3a). Neuronal activity at some electrodes raised selectively and strongly only when stimulated at the assigned location, while at others it was flatter and less specific. About 50% of this variance was explained by the specific spatial relations between ERFs and stimuli; a half-Gaussian fit to gamma responses following stimulation at each electrode’s assigned location (*R*^2^ = 0.497) shows a rapid decline of activation within less than 1.5-degree of distance between ERF center and stimulus center (Fig. 3b).

**Figure 3.**
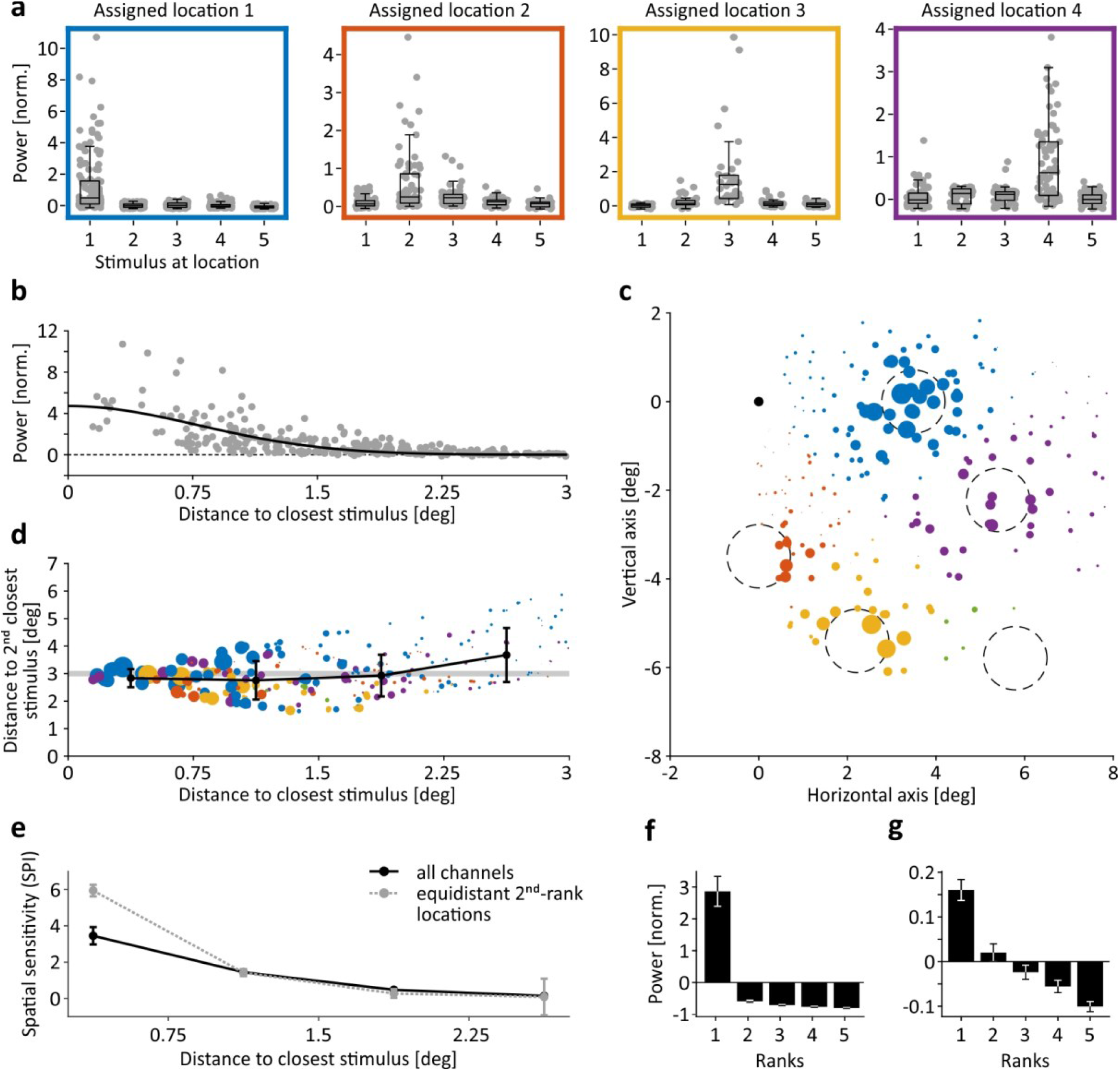
EFP modulation as function of stimulus location. (**a**) Mean gamma power (y-axes) of individual channels (gray dots) with ERFs close to stimulus location 1 to 4 (left to right) in response to stimuli shown at locations 1 to 5 (x-axes). Boxplot on top of individual channel data indicates median, 25^th^, and 75^th^percentile. Whiskers indicate interquartile range. (**b**) Mean gamma-band responses as function of distance between closest (assigned) stimulus location and ERF activation peak. Black line represents half-Gaussian fit to data. (**c**) Spatial sensitivity map in visual field coordinates. Colored dots represent individual EFP-channels, dot X-Y coordinates indicate center coordinates of the respective ERF, dot color represents assigned stimulus location, dot size is scaled by spatial sensitivity *SPI*. Dashed circles indicate size and position of large stimuli, black dot indicates fixation point. (**d**) Spatial sensitivity *SPI* as function of distance to closest and 2^nd^-closest stimulus. Conventions for size and color of dots as in (c). Black line shows mean distance of 2^nd^-closest stimulus, error bars indicate SD, gray-shaded area indicates distance to 2^nd^-closest stimulus at 3 ± 0.1-degree (equidistant 2^nd^-rank locations in (e)). (**e**) Mean spatial sensitivity as function of distance between ERF center and center of closest (assigned) stimulus. Straight black line, all channels; dashed grey line, channels with distance of 3 ± 0.1-degree to 2^nd^-closest stimulus. (**f** - **g**) Mean normalized gamma power of channels with ERF-stimulus distance below 0.75-degree (f) and between 2.25- and 3-degree (g) in response to the closest stimulus (y-axis, rank 1) and to stimuli at increasing distances (ranks 2 – 5). Error bars indicate SEM.

To investigate the spatial resolution of locational information in more detail, spatial sensitivity *SPI* was expressed as the difference Δ*WP*(*loc*)_*rnk*1,*rnk*2_ between stimuli at the closest (*rnk*1) versus second-closest (*rnk*2) location, calculated for each electrode. Because *WP* already constitutes the normalized response strength at each electrode, Δ*WP*(*loc*) preserves the amplitude of the response difference and indicates sensitivity better than indexes applying further normalization. Figure 3c shows that spatial sensitivity modulated significantly with the specific alignment of ERFs and stimulus locations, being considerably larger for ERFs in close proximity to a stimulus. Figure 3d illustrates the same data as function of distance to closest and second-closest stimulus location. Although mean distance to the second-closest stimulus was about 3-degree throughout, and mean ERF diameter was also about 3-degree, spatial sensitivity modulated clearly below this range. *SPI* decreased significantly for stimulus/ERF distances between 0.75 and 1.5-degree as compared to stimuli closer to the ERF center, and dropped to very small values for stimulus/ERF distances between 1.5 and 3-degree (Kruskal-Wallis test, *χ*^2^ = 126.29, *P* < 10^−26^, *df* = 3, pair-wise tests: all *P* < 0.011) (Fig. 3e, straight line). Difference between responses to closest and second-closest stimulus was significant at all distances (Wilcoxon signed rank tests, all *Z* > 5.01, *P* < 10^−6^), but statistical effect size *ω*^2^ decreased continuously with increasing distance between ERF center and closest stimulus location. Effect size was large^20^ only at distances below 0.75-degree (*ω*^2^ = 0.175, Fig. 3f), while medium at distances up to 1.5 and 2.25-degree (*ω*^2^ = [0.091, 0.096]) and small for distances up to 3-degree (*ω*^2^ = 0.058, Fig. 3g). Thus, locational information of gamma-band EFP responses modulates strongly within 1.5-degree distance from the ERF center, providing considerably better spatial resolution than suggested by average ERF size. We further controlled this result by considering only electrodes for which the second-closest stimulus was exactly at 3 ± 0.1-degree distance (*N* = 37), which excludes any hidden bias in Δ*WP*(*loc*) potentially resulting from a stronger influence of responses to the second-closest stimulus. For these data, the decline in *SPI* within 1.5-degree ERF/stimulus distance was even more pronounced (Fig. 3e, dashed grey line).

### Size sensitivity

Because of their fine spatial scaling, EFPs may preserve information about other, non-spatial stimulus features. First, we asked whether EFPs will be modulated by differences in stimulus size if sizes are kept in the range of ERF radii. Figures 4a - d illustrate two extremes of ERF-stimulus coverage, taken from M2. First, ERFs may perfectly overlap with stimulus coordinates such that stimuli of either size will cover and activate the ERF center region and may elicit rather similar visual responses (Fig. 4a). Yet, gamma power depended strongly on stimulus size, with the mean amplitude of large stimuli (1.4-degree in diameter) being about a factor of 1.8 and 3.5 larger than the amplitude of medium (1.2-degree) and small (1-degree) stimuli, respectively (Fig. 4b). Second, ERFs may not at all overlap with stimulus locations such that stimuli of neither size are expected to significantly modulate the visual response at the electrode (Fig. 4c). However, we still found that gamma power clearly modulated as a function of stimulus size (Fig. 4d), indicating that the effective response region measured at this electrode was apparently larger than suggested by ERF boundaries. On the one hand, this is underlining that EFPs modulate with high spatial resolution, while on the other, it is showing that EFPs may integrate over neuronal sources from well beyond the ERF boundaries (as determined by statistical response thresholds using mapping stimuli).

**Figure 4.**
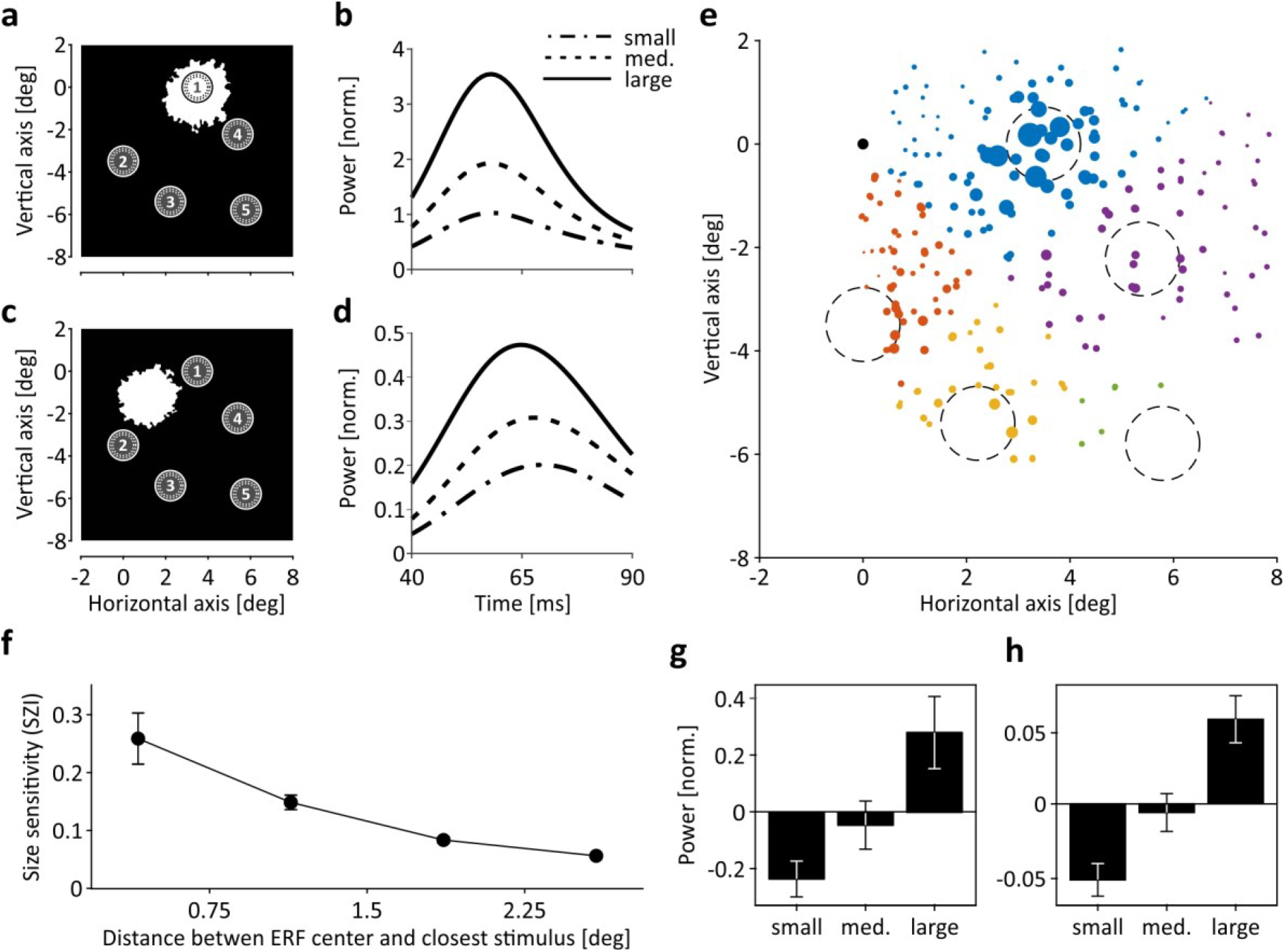
EFP modulation as function of stimulus size. (**a**) Example ERF fully overlapping with stimulus location 1. Closed circles indicate area covered by largest stimuli (1.4-degree in diameter). Dashed circles indicate area covered by medium (1.2-degree) and small (1-degree) stimuli. (**b**) Time course of mean power in ROC-selected frequency range in response to large, medium, and small stimuli. (**c**) Example ERF not overlapping with any of the stimulus locations 1 – 5. (**d**) Time course of the gamma-band response measured at this electrode. (**e**) Size sensitivity map in visual field coordinates. Colors indicate assigned location per electrode, given by smallest Euclidian distance. Size sensitivity is indicated by diameter of colored scatter and is calculated by (|Δ*WP*(*sz*)_*l,m*_| + |Δ*WP*(*sz*)_*m,s*_|)/2, with *WP*(*sz*) indicating averaged power in the selected time-frequency range over all conditions of a given size, and *l, m*, and *s* indicating large, medium, and small stimuli, respectively. Circles indicate areas covered by large stimuli. (**f**) Averaged size sensitivity as function of center-to-center distance between ERF and closest stimulus. (**g** - **h**) Averaged response strength to small, medium, and large stimuli at electrodes with ERF centers <0.75-degree distance (g) and between 2.25 and 3-degree distance (h) from stimulus center. Error bars indicate SEM throughout.

This conclusion is supported by expressing size sensitivity *SZI* as function of distance between ERF centers and closest stimulus locations (Fig. 4e), in analogy to the analysis of spatial sensitivity. Per electrode, *SZI* was calculated as the mean of the absolute differences between responses to large and medium stimuli and responses to medium and small stimuli (|Δ*WP*(*sz*)_*l,m*_| + |Δ*WP*(*sz*)_*m,s*_|)/2, with all responses being averaged across ROC-selected time-frequency bins and trials per size condition. Responses at 156 (89%) and 98 (71.5%) electrodes in M1 and M2, respectively, showed a significant modulation by size (all *P* < 0.05) and relative response differences at electrodes with larger distance to the ERF (Fig. 4h) were similar to those at close distance (Fig. 4g), with the effect size being large at all distances (*ω*^2^ = [0.19, 0.25, 0.19, 0.16]). Yet, the overall modulation strength depended significantly on stimulus/ERF center distance (Kruskal-Wallis test, *χ*^2^ = 100.28, *P* < 10^−20^, *df* = 3, pair-wise tests: all *P* < 0.025) (Fig. 4f), suggesting that EFP magnitudes modulate as a function of stimulus size on continuous spatial scales, as illustrated by the two examples shown earlier.

### Shape sensitivity

Because EFPs represent the integrated activity of presumably several cortical hypercolumns, they are unlikely to be strongly orientation-selective. EFPs may, however, possess a bias towards one orientation, depending on the specific composition of orientation columns contributing to the signal. Based on data from the ERF mapping procedure, we estimated orientation sensitivity by a method testing for statistical reliability of orientation-dependent responses^21^. In M1 and M2, we found 168 (95.5%) and 67 (48.9%) channels, respectively, exhibiting a significant bias towards one orientation, albeit absolute orientation selectivity was small, with a median orientation index *OI* of 0.115 (Fig. 5a, b). Yet, like responses to oriented bars, the response to different shapes is likely to depend, at least to some extent, on the specific composition of orientation columns activated by the stimulus. Figures 5c, d show the response of the two electrodes shown in Fig. 4a, c to each of the five shapes used for visual stimulation. Per size, all shapes were composed of the same number of pixels and were all exactly fitting into an orbit of the specified diameter. For both electrodes - the one with full stimulus/ERF overlap and the one with no stimulus/ERF overlap – gamma-band responses co-varied with the overall orientation statistics of the different shapes but not with their global orientation: The two triangular shapes (shifted by 180-degree) elicited a response with very similar time course, as did the two quadrangular shapes (shifted by 90-degree), while the response to the circle was different from all others. This pattern was found for the vast majority of channels from both animals and was evident in the averaged response over all channels (Fig. 5e). Statistical testing over all channels provided a main effect of shape (Kruskal-Wallis test, *χ*^2^ = 148.89, *P* < 10^−30^, *df* = 4), while post-hoc tests revealed no difference for comparing between responses to the two triangular (*P* = 0.99) and the two quadrangular (*P* = 0.91) shapes, but highly significant response differences (all *P* < 0.0014) for all other pair-wise comparisons. Accordingly, 138 (78.4%) and 118 (90.1%) individual channels of M1 and M2, respectively, were significantly modulated by angularity (Kruskal-Wallis tests, all P < 0.05). At 137 of these channels (53.4%), two of the three angularity conditions were significantly different, and at 12 channels (4.7%) there was a significant response difference between all three angularity conditions (pair-wise tests with Tukey-Kramer multiple comparison correction, all P < 0.05).

**Figure 5.**
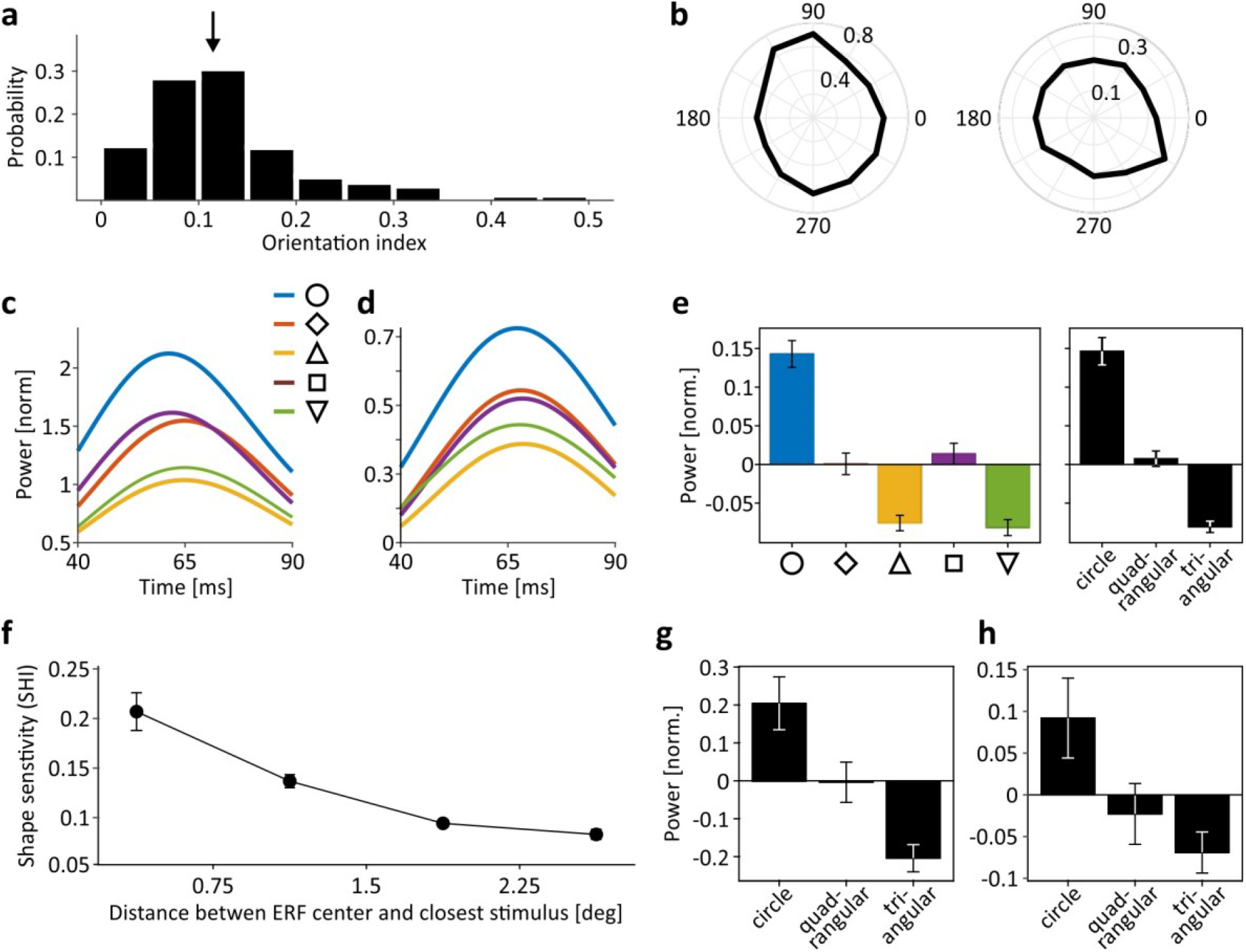
EFP modulation as function of stimulus orientation and shape. (**a**) Distribution of orientation indices from all channels with significant orientation bias (*N* = 235). Arrow indicates median (*OI* = 0.115). (**b**) Two example electrodes exhibiting orientation bias close to median (left) and close to max (right) *OI*. Numbers on horizontal and vertical axes, stimulus orientation; numbers within polar plots, normalized response strength. (**c** - **d**) Time course of responses to different shape conditions of an example channel with full overlap between ERF and stimulus area (c) and another channel with no overlap (d). Channels are the same as shown in Fig. 4a, c. (**e**) Averaged responses over all channels to each of the five shape conditions (left) and after merging conditions with identical angularity (right). (**f**) Averaged shape sensitivity as function of distance between ERF center and closest stimulus. Shape sensitivity is given by (|Δ*WP*(*sh*)_*c,q*_| + |Δ*WP*(*sh*)_*q,t*_|)/2, with *WP*(*sh*) indicating averaged power in the selected time-frequency range over all conditions of given angularity, and *c, q*, and *t* indicating responses to circular, quadrangular, and triangular stimuli, respectively. (**g** - **h**) Averaged response strength to circular, quadrangular, and triangular stimuli at electrodes with ERF centers <0.75-degree distance (g) and between 2.25 and 3-degree distance (h) from stimulus center. Error bars indicate SEM throughout.

Like spatial and size sensitivity, shape sensitivity modulated significantly with the distance between ERF and stimulus center (Kruskal-Wallis test, *χ*^2^ = 81.09, *P* < 10^−16^, *df* = 3) (Fig. 5f). Shape sensitivity was expressed as the mean of the absolute differences between responses to circular and quadrangular shapes and responses to quadrangular and triangular shapes (|Δ*WP*(*sh*)_*c,q*_| + |Δ*WP*(*sh*)_*q,t*_|)/2. It was largest at ERF/stimulus distances below 0.75-degree (Fig. 5g) and decreased significantly for distances up to 2.25-degree (pair-wise tests: all *P* < 0.022), and was still present at distances up to 3-degree (*P* < 10^−6^) (Fig. 5h). Effect size was large at all distances (all *ω*^2^ > 0.1812). Hence, in summary, EFPs keep significant information about object shape as expressed in overall orientation statistics, independent of other stimulus parameters contributing to the signal.

### Color sensitivity

Two channels of double-opponent cells in blob regions and complex-equiluminant cells outside blobs contribute to color processing in V1^14,22,23,24,25^. Though the number of neurons showing a chromaticity-dependent response bias was recently suggested to be significantly larger than previously assumed^26^, color preference in V1 is usually weak and highly variable among neurons^27^. As such, and because V1 does not contain clusters with identical color preference that would predict chromaticity-specific responses for the set of equiluminant stimuli we used, differences in color were not primarily expected to induce significant color-specific EFP modulation. In line with this assumption, the ROC analysis to find most-informative time-frequency bins for a given stimulus feature did not indicate significant detection rates during the early response from 40 to 90 ms post-stimulus onset, as it did for location, size, and shape. Contrary to this assumption, however, it did indicate significant detection rates during a later response window, 120 to 175 ms post onset (Fig. 2). Figure 6a provides an example of the response time course at a single electrode of M2. The early peak in gamma power showed some, yet small, difference in response to differently colored stimuli, whereas during the later response, red and blue stimuli elicited obviously higher activation than other stimuli. Figure 6b shows another single channel, taken from M1, with basically the same response pattern and again, stronger activation in response to red and blue stimuli during the later period. Interestingly, this color-related response bias did not vary over electrodes and was basically the same at all electrodes. On average, blue and red stimuli induced around 10% more power during the early response, and around 90% during the later response (Fig. 6c). Statistically, this was equivalent to only 3, and as much as 210, of 313 channels displaying a significant, color-dependent response difference during the early and late response period, respectively (Kruskal-Wallis tests, *P* < 0.05, *df* = 4). To further test whether the difference in statistical outcome was due to differences in overall activation between early and late response epochs, we computed a color selectivity index *CSI* by first, subtracting the response to the color eliciting smallest gamma modulation from the response to all other colors and subsequently, taking the sum of the absolute differences between mean responses to different colors, divided by the overall mean. *CSI* yields values ≥ 0 which are independent of absolute activation and allows, therefore, to investigate color selectivity during the initial onset response with later response epochs. We found that *CSI* values were significantly larger during the later epoch (Wilcoxon signed rank test, *Z =* 10.69, *P* < 10^−25^, *N* = 313), indicating that the response bias towards red and blue indeed increased over time and was unlikely to result from a difference in luminance (Fig. 6d). Rather, it is in line with recent studies showing general differences in hue-specific LFP gamma power modulation^19,28,29^ and indicates that the EFP reliably captures such hue-dependent differences.

**Figure 6.**
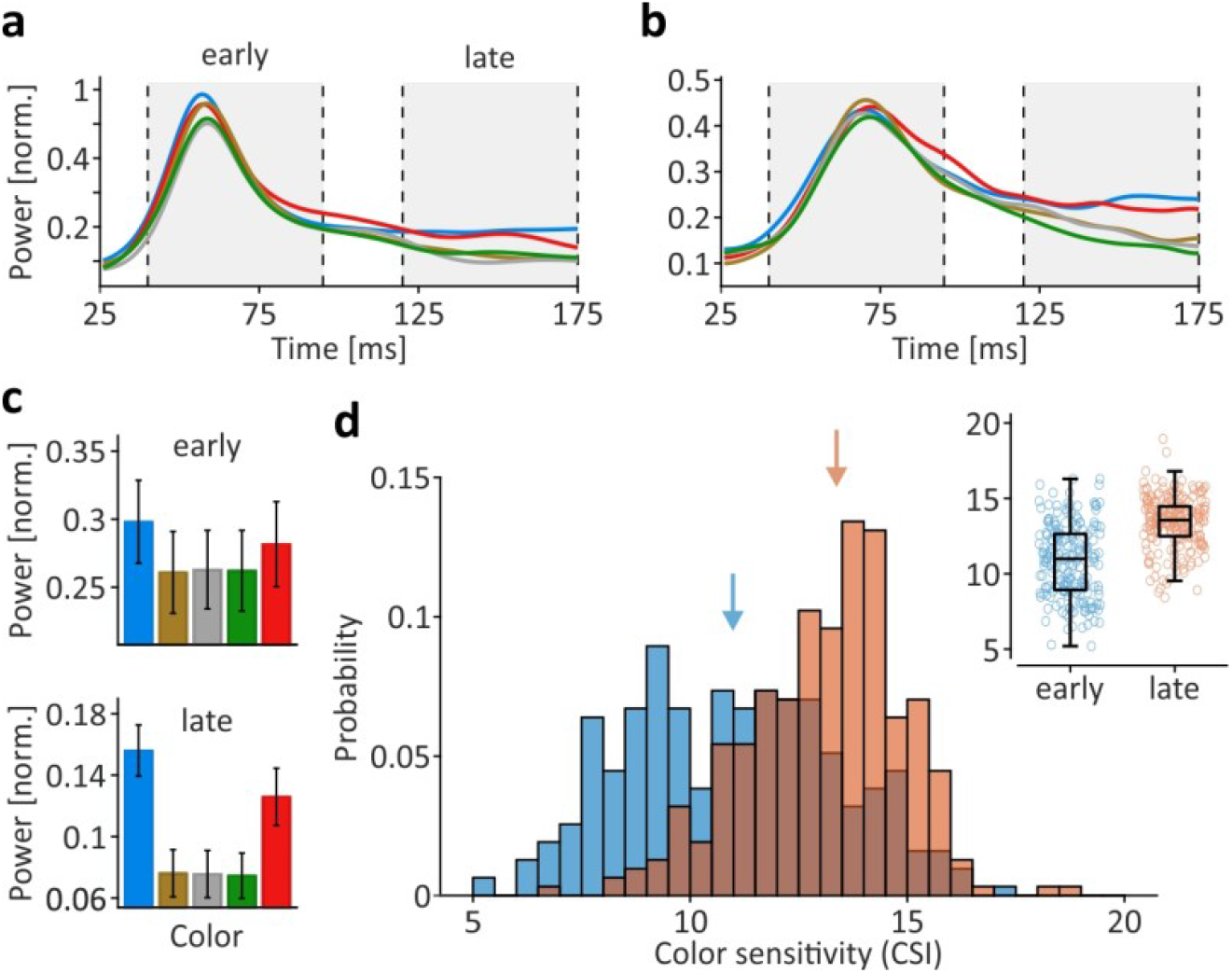
EFP modulation as function of color. (**a** - **b**) Time course of mean power in ROC-selected frequency range (a: M2, 47 – 96 Hz; b: M1, 42 – 145 Hz) in response to differently colored stimuli. Line color indicates stimulus color. Grey-shaded areas indicate response periods during transient stimulus onset (‘early’) and sustained (‘late’) activation. (**c**) Power in ROC-selected time-frequency range, averaged over all electrodes, during early and late response periods. Bar color indicates stimulus color. Error bars, SEM. (**d**) Color index distribution during early (blue) and late (orange) response periods. Color index *CSI* is independent of absolute activation. Arrows indicate median *CSI* during early and late responses. Inset shows *CSI* of individual electrodes, boxes indicate 25^th^ and 75^th^percentile including median, whiskers indicate interquartile range.

### Single-trial classification precision

The results shown so far indicate that non-spatial stimulus features induce characteristic modulations of the EFP gamma power, which were qualitatively similar at all electrodes. We therefore finally asked to what extent pre-knowledge about these characteristics allows to distinguish single trials from objects shown at the same location. Specifically, because size, shape, and color of a stimulus may be chosen to elicit either low, medium, or high gamma power on average, we asked whether this modulation is sufficiently invariant to allow support vector machines to classify single trials solely based on non-spatial stimulus features. First, responses to a combination of stimulus features presumably eliciting high gamma power (large, blue, circle) were tested against a combination of features presumably eliciting low gamma power (small, green, triangle), using the trials in response to a stimulus at each electrode’s assigned location. Trials were collapsed into three numbers, each number representing the trial’s maximal power value in the respective time-frequency range of size, shape, and color. We found that 122 channels were performing significantly (*P* < 0.01) above chance level and of these, 62 had detection rates of more than 70% correct, and many were performing better than 80%. Likewise, for testing another high vs. low gamma combination (large red circle vs. small brown inverse triangle), 113 channels were found to perform significantly above chance level, and 57 channels had detection rates of over 70% correct (Fig. 7a, red lines). We also tested objects eliciting high or medium-sized gamma responses against objects eliciting medium-sized or small responses, respectively, and still found a significant number of electrodes performing above chance level, some reaching detection rates around, or even clearly above, 80% correct (Fig. 7a, blue and green lines). Figure 7b shows that classification performance depended significantly on the center-to-center distance between ERF and assigned stimulus location (rank correlation, *R* = −0.652, *P* < 10^−16^) and was best for stimuli in close proximity to the ERF center. For five electrodes of both M1 and M2, with ERFs at location 1, we tested all possible object combinations for the high vs. low gamma comparison (large red (blue) circle vs. small green (gray, brown) triangle (inverse triangle); *N* = 12). Despite the fact that some ERFs had only partial overlap with the stimulus location (Fig. 7c), average classification performance in monkeys M1 and M2 was as high as 75.6% ± 4.6% SD and 88.5% ± 1.5% SD, respectively (Fig. 7d). Thus, although EFPs represent the integrated activity over presumably tens of thousands of neurons, they depict slight differences in non-spatial visual features that are not only visible after averaging over many trials but allow reliable classification of single trials reduced to as few as three numbers.

**Figure 7.**
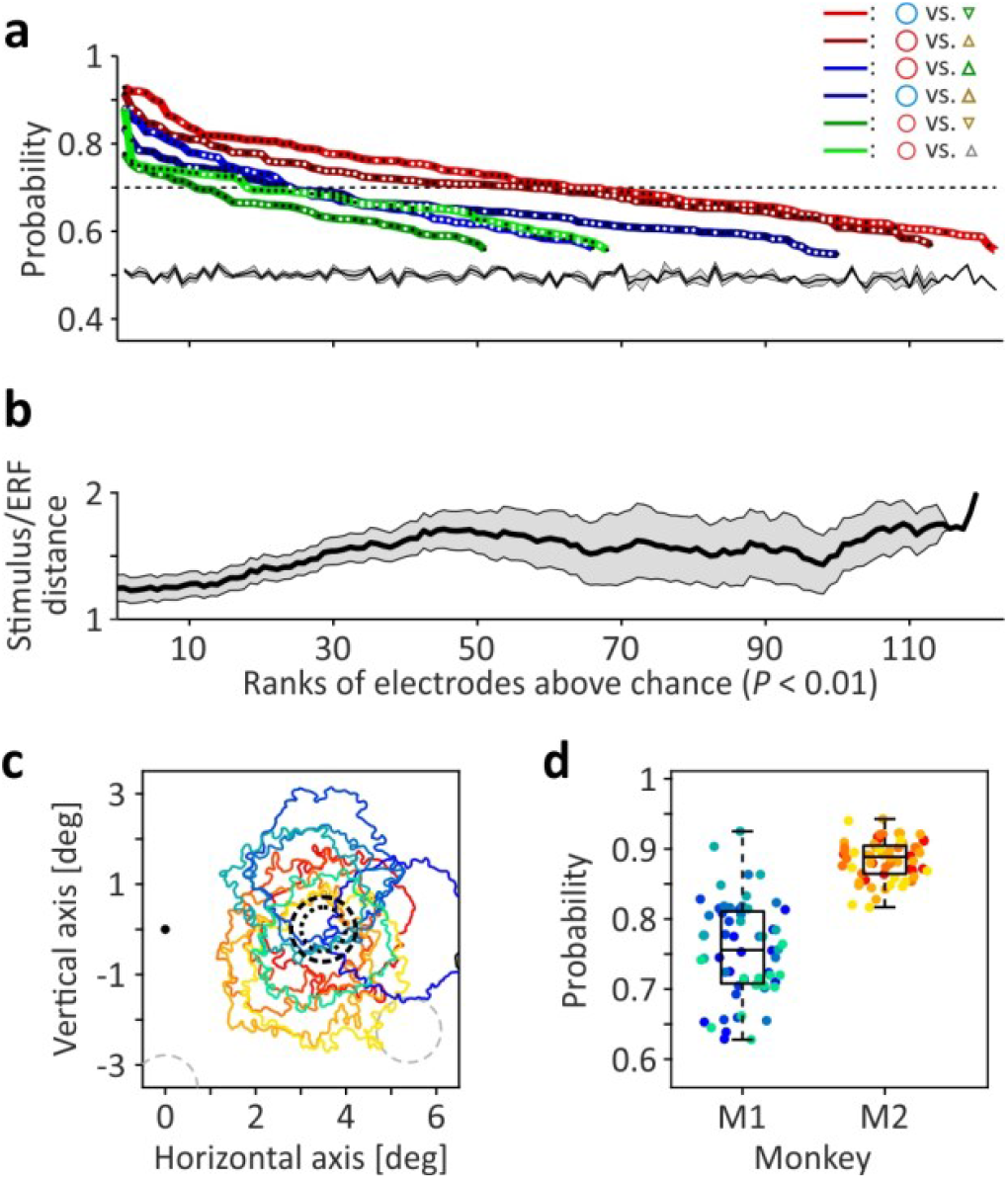
Single-trial object classification based on non-spatial features. (**a**) Support-vector machine performance at individual electrodes for six different classification conditions, comparing objects eliciting high vs. low gamma (red lines), high vs. medium gamma (blue lines), and medium vs. low gamma (green lines). Black and white dots plotted on lines indicate individual electrodes of M1 and M2, respectively. Only electrodes with performance significantly (*P* < 0.01) above chance level are shown, ranked by classification rate. Experimentally determined mean chance level ± SEM is given by black line and grey-shaded area. (**b**) Mean center-to-center distance between ERF and closest stimulus of electrodes with significant classification performance. Black line indicates moving average over 25 electrodes, shifted by 1 electrode, grey-shaded area indicates corresponding SEM. (**c**) ERF outlines of five selected electrodes per monkey (M1: blueish/greenish, M2: reddish/yellowish). (**d**) Single-trial classification performance of electrodes shown in (b) to each of 12 high vs. low gamma conditions. Boxes indicate 25^th^ and 75^th^percentile including median, whiskers indicate interquartile range.

## Discussion

Brain signals for both clinical approaches and basic research may be acquired in different ways, ranging from electroencephalography (EEG) over intracranial recordings from the brain’s surface to multi- and single-unit recordings at the macroscopic, mesoscopic, and microscopic level, respectively. Intracranial surface recordings as a mesoscale signal have gained much interest in recent years; for clinical settings, they offer a much better signal-to-noise ratio than EEG and for basic research, they allow to study dynamic interactions simultaneously over large and/or distant brain regions. In both clinical settings and basic research, they are most widely implemented by subdural implantation of electrodes (referred to as electrocorticography, ECoG). Yet, placement of electrodes below the dura introduces the risk of peri- and post-operative complications and likely limits the duration over which signals can be recorded, particularly in clinical settings. An alternative to opening the dura is the placement of electrodes on top of it and recording epidural instead of subdural field potentials. Although attractive from the perspective of safetiness, only few studies have performed EFP recordings and little is known about their basic properties and functional specificity, making them, in turn, still a rare choice.

We here make use of the high spatial resolution and functional specificity of monkey V1 to investigate EFP sensitivity to spatial and non-spatial visual stimulus features, and show that EFPs modulate on much finer spatial scales than suggested by the size of their receptive fields, and moreover, that modulation caused by non-spatial features is highly specific and allows very good single trial classification rates. EFPs, hence, constitute a highly selective signal, both in terms of locational and functional specificity, that is a valid alternative as source of information on distributed cortical activity.

### Functional specificity of epidural field potentials

EFPs mainly arise from electrical activity in superficial layers of the cortex^30^. Most of the apical dendrites there come from layer 5 pyramidal neurons, which, in a simplified view, can be conceived as a parallel arrangement of sinks and sources (that is, dipoles) that would produce the electrical field recorded by the electrode (although dendritic morphology and temporal dynamics are complex and primary sources may also include mono- and multipolar components^31^). Because the amplitude of a given dipole is smaller the larger the distance to the recording site, the EFP represents a spatially weighted superposition of potentials from spatially distributed sources. As a consequence, its amplitude, shape, and spatial spread is primarily dependent on the number and magnitude of, and the synchrony among, the potentials of the contributing neurons^32,33,34^.

The average size of epidural RFs was recently estimated to be around two to three times the size of intracortically recorded LFP-RFs when mapped with the same protocol^11^ and using corresponding statistical thresholds^12^ for specifying their boundaries, and well in the range of subdurally recorded field potentials^6,35^. The results of the current study suggest an even larger effective size of EFP-RF’s, as indicated by the small, yet significant location and size sensitivity at stimulus coordinates in the periphery of the ERF (Fig. 3g and 4h) and at locations well outside ERF boundaries (e.g. Fig. 4c, d). A likely reason for this is the difference in the stimuli used: Because the extent of spatial spread is strongly dependent on the coherence among the contributing signals^34^, the flashing of visual objects as performed in the current study is expected to induce stronger synchronization during the initial stimulus onset response of the neurons than the moving bar stimuli used for mapping, which slowly enter the RF.

While this relatively far-reaching spatial spread could be taken as indicator for poor spatial sensitivity, we found that EFPs in fact modulate on much finer scales. This is best illustrated by the results on size sensitivity: Small, medium and large stimuli each differed in size by only 0.2-degree diameter, yet stimulus sizes were associated with significantly different gamma power at all stimulus-ERF distances. Given a number of about 2.5 * 10^5^neurons under 1 mm^2^ monkey V1 cortex^36,37^, even small changes in stimulus size are expected to significantly change the number of neurons carrying the EFP and eventually induce significant differences in gamma power, which finally gives rise to strong and consistent size sensitivity as observed at the vast majority of array electrodes. Similar findings on size sensitivity have previously been obtained for the LFP^38,39,40,41^ and for subdurally recorded ECoG^42^, although stimulus size differences were considerably larger in these studies as compared to ours.

Furthermore, we found a strong dependence of EFP gamma power on stimulus shape, given by reliably different amplitudes for objects of different geometrical shape but no influence of global stimulus orientation. Because LFP gamma power is known to express some degree of orientation sensitivity when tested with gratings^40,43^, a possible reason is a shape-specific activation of spatially distributed clusters of orientation-selective neurons. In support of this, we found a slight yet significant orientation bias in EFP responses, in line with a recent investigation of subdural ECoG^42^. An alternative hypothesis, however, not relying on orientation-selectivity, is a general difference in the number of neurons activated by the three geometrical bodies: Regardless of whether size is defined by the diameter of the surrounding orbit (as we did) or by object surface area (as an alternative), different shapes will either cover a differently sized area of the ERF (in the former case) or extend differently far into its periphery (in the latter). Thus, distinct geometrical bodies will activate unique pools of spatially distributed neurons, and, consequently, the weighted superposition of these neurons’ potentials will be specific for each of the shapes. Orientation-selective responses from local clusters of neurons, as discussed above, may contribute to this. In conclusion then, size sensitivity in V1 EFPs would emerge as a secondary property caused by stimulus-specific differences in spatio-temporal activation patterns of large neuronal populations. Note, however, that recent work reported shape selectivity also as a primary property of V1 population activity, yet using two-photon imaging, which has significantly higher spatial resolution^44^.

Finally, red and blue stimuli elicited stronger gamma responses, albeit during a later period after the initial onset response, as compared to both equiluminant green and brown color stimuli and achromatic stimuli. This result is unlikely to be due to columnar organization within V1; firstly, hue maps in V1 (basically overlapping with blob regions) have a diameter of about 160 µm and contain the full gamut of colors^45^, such that recordings of color-selective responses requires microscale spatial resolution. Secondly, the bias towards red and blue stimuli was evident at all electrodes likewise, indicating some general differences in processing of short- and long-wavelength hues as compared to mid-wavelength hues. Stronger gamma power to bluish and reddish hues has also been found in a recent study investigating hue-specific differences in LFP gamma power^28^. Yet, these authors also found generally higher power for chromatic over achromatic stimuli, using full-screen stimulation, a pattern that was not evident with the small objects on dark background we applied in our study. Such small differences in stimulation were recently shown to have a significant impact on color-specific gamma modulation^29^, and may explain some differences between our study and the studies of others. Independent of the specific modulation, the results show that the EFP is reliably catching up color-specific information which could then be used for single-trial classification, as we did for distinguishing objects shown at the same location.

### Time-frequency ranges for distinct stimulus features

Several studies investigating quantitatively LFP and ECoG spectral composition reported distinct frequency ranges for different stimulus features^17,18,19^. We applied receiver-operating characteristics to isolate most-informative time-frequency ranges and found locational sensitivity mainly in the high gamma range (> 85 Hz), compatible with elevated synaptic and spiking activity^32,41,46^ as a result of better stimulus placement relative to ERF coordinates. Size and shape sensitivity elicited prominent gamma increases in mid-frequency range (30 – 80 Hz) during the initial onset response, with minor differences between monkeys. Corresponding results were reported by previous studies investigating LFP modulations as function of stimulus size and orientation^38,39,40,42,43^. Because shape sensitivity is most likely resulting from stimulus-dependent differences in effective RF coverage and orientation statistics, the matching of frequency-ranges for size- and shape-induced modulation is consistent. Color-specific differences had maxima in two gamma frequency ranges in M1, a higher (80 – 145 Hz) and a lower (42 – 58 Hz) one, while only the lower showed up in M2. These modulations were delayed in time as compared to the former features, and had lower amplitude. Thus, the ROC-procedure revealed strongest modulations in two distinct gamma frequency ranges, consistent with earlier reports^41^, with the high-gamma presumably reflecting increased spiking activity and the lower indicating induced oscillatory rhythms, which depend on both stimulus properties and cognitive processes^47^. We did not attempt to investigate whether different stimulus properties may have power maxima in slightly shifted frequency ranges, as described for stimulus contrast variations^18,19,48^. The functional impact of such frequency shifts is a matter of debate, with some arguing it is indicating a limitation for neuronal communication^48^ and others arguing for the opposite, more reliable phase coordination by stronger synchrony through non-stationary frequency modulations^19,49^.

## Conclusion

The results of the current study provide evidence that the EFP is holding highly selective information about constituent features of activating stimuli and in particular, that non-spatial stimulus features modulate the EFP in a highly consistent manner. This allows to purposefully use these features for increasing decoding rates in brain-computer interfacing and clinical settings. Moreover, because data were obtained from multiple recording sessions spread over several weeks, the results are rather underestimating the sensitivity of the EFP than overestimating, which may still be significantly increased by keeping session-wise variability at low level. This opens new perspectives for less traumatic implantation of arrays and for their long-term application in both clinical settings and basic research on meso- and large-scale network interactions and also points on the importance of future research on technological improvements of epidural multielectrode arrays.

## Methods

### Subjects and ethical statement

Recordings were performed in two male macaque monkeys, M1 (13 yr., 12 kg) and M2 (14 yr., 11.5 kg). M1 was housed pairwise in a large indoor compartment with daily access to an equally sized outdoor compartment. M2 was housed in an indoor compartment with visual and auditory contact to other monkeys. All compartments were enriched by a manifold of toys, puzzles, and climbing opportunities. Both monkeys were familiar with standard laboratory procedures. They were given water and/or fruit juice during training and recording sessions as reinforcer for performing a dimming task at fixation. On non-working days, they received water and varieties of fruits in their home compartment. Primate food pellets and seeds were offered all days. Both monkeys were implanted with an epidural multielectrode array covering the left occipital hemisphere and extending towards the lunate sulcus. Technical details of the array^16^ and its implantation^11^ are given elsewhere. All surgeries were performed under initial ketamine/medetomidine anesthesia supplemented with isoflurane and, if necessary, maintained by propofol/remifentanil, and under strictly aseptic conditions. Postoperative analgesia was provided by Caprofen. Health and well-being of the animals were checked by daily visual inspection, frequent control of body weight, and regular veterinarian assessment. All experimental, surgical, and behavioral procedures followed the Directive 2010/63, issued by the European Commission, and the Regulation for the Welfare of Experimental Animals, issued by the Federal Government of Germany, and were approved by the local authorities.

### Visual stimulation

Visual stimuli (Fig. 1B) were shown on a CRT display (1.152 x 864 px resolution, 100 Hz refresh rate) and consisted of five different shapes (circle, diamond, triangle, square, and inverse triangle) in three different sizes. Size was defined by an orbit of 1.4-degree (large), 1.2-degree (medium) and 1.0-degree (small) diameter (Fig. 1A), into which objects had to fit exactly. Objects of same size were matched in number of pixels. All objects were shown in four equiluminant colors (blue, brown, green, and red) and one achromatic (gray) color, with isoluminance (10 cd/m^2^) determined physically by repeated measurements, and displayed on a dark (∼0 cd/m^2^) background. Objects were presented at five different locations at 3.5, 5.8, and 8.2-degree eccentricity. Object locations (Fig. 1a) were chosen to provide different degrees of coverage with epidural receptive fields, to allow testing EFP modulations as function of density and distance. Combination of 5 shapes x 3 sizes x 5 colors x 5 locations provide a total of 375 stimulus conditions.

Stimuli were shown during a simple dimming task requiring the monkeys to indicate a slight luminance change of the fixation point (0.2 x 0.2-degree). Monkeys initiated a trial by gazing at the fixation point and pressing a lever. 750 ms after trial initiation, three visual objects were presented sequentially, each for 800 ms (Fig. 1c). Consecutive stimuli were separated by 250 ms blank periods and never consisted of the same condition. Dimming of the fixation point occurred at a pseudorandom point in time within maximally 1.1 sec after offset of the third stimulus in a sequence. Monkeys were rewarded for indicating detection of the luminance change within a 750 ms response window, starting 200 ms after dimming. Trials were terminated without reward if the monkey responded too soon or too late, or if the monkey’s gaze deviated by more than ∼1-degree from the fixation point.

Epidural receptive fields (ERFs) were determined by an automated mapping procedure using oriented grey bars (size: 0.24 x 23.8-degree) moving along a 19.1 -degree trajectory in one of 12 directions, spaced by 30-degree and centered at x = 2.4-degree, y = -3.6-degree. Response maps were obtained by back-projecting wavelet-transformed epidural responses in broadband gamma frequency range (60 – 150 Hz) to actual stimulus location. Details of the method are given elsewhere^11,12^. Receptive fields were defined as area of ≥ 1 square-degree size and *Z*-scores ≥ 1, identical to earlier work^11^. Orientation index *OI* was calculated as Michelson contrast between responses to the most activating bar orientation *θ* and an orientation 90-degree away from that:

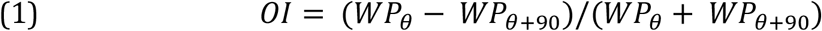

During mapping, monkeys performed a dimming task at fixation.

### Data acquisition and preprocessing

Epidural arrays^11,16^ consisted of 202 hexagonally arranged recording electrodes, each 560 µm in diameter, with 1.8 mm center-to-center distance. EFPs were recorded at 25 kHz sampling rate using recording equipment from Multichannel Systems (Reutlingen, Germany; recording chain: MPA 32, Sc2×32, PGA64, USB-ME256). Signals were referenced against a large reference electrode on top of the array, facing the skull. Additional ground electrodes were implanted over right frontoparietal cortex. The monkeys’ eye position was recorded using a custom-made video-oculography system with 0.2-degree spatial resolution. Raw EFPs, analog eye signals and a 50 Hz socket signal were stored for offline analysis. To prevent over-estimation of EFP selectivity, data were recorded within 13 and 25 sessions over a time period of 17 and 16 weeks in M1 and M2, respectively, thus likely showing higher trial-by-trial variability than data from a single session and providing a robust estimate of EFP selectivity over time.

All data was low-passed filtered (< 300 Hz, finite impulse response filter in forward and backward direction, cutoff at 150 Hz) and down-sampled to 1 kHz. 50 Hz noise from the EFP was removed using the socket signal. All data was wavelet-transformed using Morlet wavelets, as described in detail elsewhere^11^. Single-trial wavelet power was baseline-normalized by subtracting the frequency-wise mean power during the 500 ms baseline period at trial beginning, and then dividing through it.

### Electrode selection and trial rejection

Only EFPs originating from neuronal populations in V1 were used for analysis. Electrodes located anterior to the lunate sulcus and electrodes not delivering a significantly modulated ERF were excluded from analysis. The final database consisted of 178 and 137 electrodes from M1 and M2, respectively.

Data analysis was limited to the time period of 26 ms to 175 ms following stimulus onsets. Prior to analysis, responses during this interval (denoted as single trials from hereon) and during the interval used for baseline correction (201 – 700 ms post-trial start) were checked for artifactual activity patterns by a semi-automatic procedure. First, trials with temporally no or sparse modulation were rejected; second, trials with power above or below the mean broadband power (30 – 160 Hz) ± 4 SD of all (remaining) trials of a given stimulus condition and electrode were rejected; third, trials with highest mean broadband power (above 80^th^percentile of all (remaining) trials of a given stimulus condition at majority of electrodes) were tested for unusual strong correlations across electrodes by applying Pearson correlation for each stimulus condition. Trials were rejected if the mean over all pair-wise correlation coefficients was ≥ 0.5. If necessary, additional trials with suspiciously high power were excluded by final visual inspection.

### Receiver-operating characteristics

Wavelet-transformed EFPs of each monkey were first down-sampled to time bins of 5 ms length and were then subjected to receiver operating characteristics (ROC) analysis to identify most-informative time-frequency ranges. Per electrode and time-frequency bin, all data from trials belonging to one stimulus condition were tested against the corresponding data from all trials of the remaining stimulus conditions of the same feature category (i.e. size ‘large’, 1.4-degree diameter vs. size ‘medium’ and ‘small’, 1.2 and 1.0-degree diameter), thus providing the area-under-the-ROC curve (AUC) per stimulus condition, electrode, and time-frequency bin. AUC values of all electrodes and conditions were used to construct grand-variance maps expressing the *Z*-transformed variance over all AUC values in time-frequency space (Fig. 2). Variance is highest in the time-frequency range where a given stimulus condition is well separable from other conditions at many electrodes, while small if separability is poor and AUC values are all close to 0.5. A detailed, step-by-step description of the procedure is given elsewhere^11^. For each of the four categories, analysis was based on the most informative time-frequency range (white rectangles in Fig. 2).

### Feature sensitivity computation

EFP sensitivity to features of the four categories location, size, shape, and color were analyzed by averaging over the time-frequency bins of the respective category at each electrode. Per category, all trials were considered. Spatial sensitivity *SPI* of each electrode was computed by comparing mean power *WP* in response to stimulation at each of the five stimulus locations and calculating the absolute difference in power between locations eliciting the highest (*rnk*1) and the second-highest response (*rnk*2):

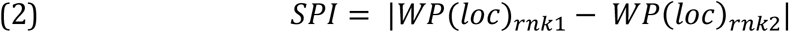

Size sensitivity *SZI* was calculated as the mean of the absolute difference in power between responses to large (*l*) and medium (*m*) stimuli and responses to medium and small (*s*) stimuli:

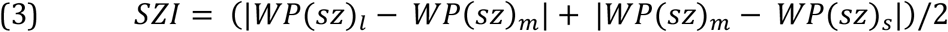

Shape sensitivity *SHI* was calculated accordingly, using mean *WP* in response to circular (*c*), quadrangular (*q*), and triangular (*t*) shapes:

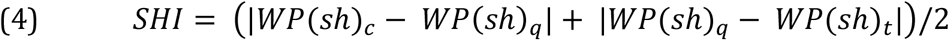

Indexes *SPI, SZI*, and *SHI* are larger the larger the absolute difference in *WP*, i.e. they scale with overall activation. In contrast, color sensitivity *CSI* was computed to be independent of overall activation, to test for differences in *CSI* during the course of a trial, particularly between transient and more sustained response periods. *CSI* was calculated by first, taking the mean responses to the five colors *WP*(*col*) and subtract the minimum of these from each, and second, using these corrected *cWP* values for calculating the sum of the absolute differences between responses to differently colored stimuli, divided by their overall mean:

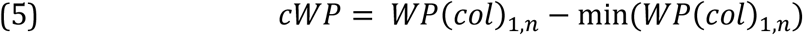

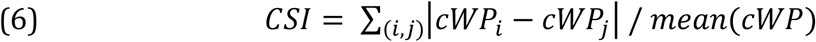

Subtraction of the minimum response and division by the overall mean provides *CSI* values independent of the overall activation.

### Support vector machines

Single-trial decoding performance was estimated using nonlinear support vector machines (SVMs) with Gaussian radial basis function kernel, as provided in the *libsvm* toolbox for Matlab^50^. Per electrode, an equal number of trials was used for the two stimulus conditions to compare. If applicable, trials were randomly drawn from the condition with the higher quantity. All trials were scaled to [0, 1]. Trials were classified using a leave-one-out procedure, i.e. each trial in the data was used once for classification, while remaining trials served as training set. SVMs were trained by first subjecting the training set to SVM classification with 5-fold cross validation for initial estimation of best values for parameters C and gamma, and subsequent training of SVMs using the optimal C and gamma values. Parameters C and gamma represent SVM parameters for the separation of data points into two classes and definition of support vectors. Each test trial was assigned to the stimulus class with highest probability. Training and classification were repeated ten times per electrode and stimulus pair. Performance of electrodes was computed by comparing the assigned stimulus class of all trials to their true stimulus class. Chance level was 50% per definition but, for statistical reasons, was confirmed experimentally by repeating the training/classification procedure with shuffled class labels, again performing ten rounds per electrode and stimulus pair.

### Experimental design and statistics

For investigating EFP sensitivity to stimulus features location, size, shape, and color, we always referred to the entire set of data, with trials sorted accordingly. All data was statistically investigated by non-parametric methods. If not stated otherwise, significance is reported at α ≤ 5%. Statistical effect size is given as omega square^11^. Classification performance of individual electrodes was tested by comparing the performance of ten full classification runs per electrode with true labels against another ten runs with shuffled labels, using Mann-Whitney U-test. Electrode performance was considered significantly above chance at α ≤ 1%.

## Acknowledgments

The work was supported by DFG grant WE 5469/3-1, Tönjes-Vagt Foundation (Project XXXI), University of Bremen (ZF grant and Creative Unit I-See), and a scholarship from the German Academic Scholarship Foundation. The authors acknowledge Iris Grothe for valuable comments on an earlier draft and Peter Bujotzek, Katja Taylor, Katrin Thoss, Ramazani Hakizimani, and Constantin Neagu for their support.

## Author Contributions

B.F. and D.W. conceived and designed research, B.F. performed experiments, B.F. and D.W. analyzed and interpreted data, D.W. prepared figures and wrote the manuscript, B.F. and D.W. approved final version of manuscript.

